# Predatory colponemids are the sister group to all other alveolates

**DOI:** 10.1101/2020.02.06.936658

**Authors:** Denis V. Tikhonenkov, Jürgen F. H. Strassert, Jan Janouškovec, Alexander P. Mylnikov, Vladimir V. Aleoshin, Fabien Burki, Patrick J. Keeling

**Author notes:** Shared first authorship, both authors contributed equally (Denis V. Tikhonenkov, Jürgen F. H. Strassert). Institute of Biology, Free University of Berlin, Königin-Luise-Straße 1–3, 14195 Berlin, Germany (Jürgen F. H. Strassert). Corresponding authors Email addresses. Jan Janouškovec, Vladimir V. Aleoshin, Fabien Burki, Patrick J. Keeling.

## Abstract

Alveolates are a major supergroup of eukaryotes encompassing more than ten thousand free-living and parasitic species, including medically, ecologically, and economically important apicomplexans, dinoflagellates, and ciliates. These three groups are among the most widespread eukaryotes on Earth, and their environmental success can be linked to unique innovations that emerged early in each group. Understanding the emergence of these well-studied and diverse groups and their innovations has relied heavily on the discovery and characterization of early-branching relatives, which allow ancestral states to be inferred with much greater confidence. Here we report the phylogenomic analyses of 313 eukaryote protein-coding genes from transcriptomes of three members of one such group, the colponemids (Colponemidia), which unambiguously support their monophyly and position as the sister lineage to all other alveolates. Colponemid-related sequences from environmental surveys and our microscopical observations show that colponemids are not common in nature, but diverse and widespread in freshwater habitats around the world. Studied colponemids possess two types of extrusive organelles (trichocysts or toxicysts) for active hunting of other unicellular eukaryotes and potentially play an important role in microbial food webs. Colponemids have generally plesiomorphic morphology and illustrate the ancestral state of Alveolata. We further discuss their importance in understanding the evolution of alveolates as well as origin of myzocytosis and plastids.

**Highlights:**

- Phylogenomics resolves Colponemidia as a sister group to all other alveolates
- The ancestor of all alveolates was a biflagellate predator feeding by phagocytosis
- Colponemids may illuminate the ancestral states of apicomplexans, dinoflagellates, and ciliates
- Colponemids are geographically widespread in freshwater habitats

## 1. Introduction

Alveolates are one of the largest major groups of eukaryotes with over ten thousand described species and much larger yet undescribed species diversity based on environmental sequence data (Chambouvet et al., 2008; López-García et al., 2001; de Vargas et al., 2015). These organisms have a special type of cell coverings consisting of a plasmalemma and alveoli — abutting single-membrane flattened sacs which probably derived from the endomembrane system and always subtend the plasma membrane (Gould et al., 2008). Most alveolates belong to one of three major groups: apicomplexans (e.g., the malaria parasite *Plasmodium*), dinoflagellates (e.g., the coral endosymbiont *Symbiodinium*), and ciliates (e.g., the model organisms *Tetrahymena* and *Paramecium*). These three groups are among the most ecologically successful eukaryotes on Earth, and are also of evolutionary importance since each lineage also displays a number of distinctive innovations. For example, the apicomplexans infect virtually all known animals and this process is mediated by a complex suite of structures called the apical complex (Katris et al., 2014). Ciliates have evolved a large cell size and structures that make them effective predators, as well as a separation of germ and soma within a single cell. Dinoflagellates have evolved a variety of unique features, including several aspects of genomic organization and function, as well as a wide range of trophic strategies including photosynthetic algae, predators, and parasites. The alveolates have also served as models to understand organelle evolution; mitochondrial genomes and respiratory chains are modified in each of the three groups in different ways (Smith et al., 2007; Nash et al., 2008), and apicomplexans and dinoflagellates (together with their closest relatives collectively known as myzozoans) contain plastids that have adopted diverse functional and genomic states (Janouškovec et al., 2017a).

Reconstructing the deep evolutionary transitions that gave rise to these three well-studied and diverse groups has relied heavily on the discovery and characterization of early-branching lineages, which allow ancestral states to be inferred with much greater confidence. Such deep-branching lineages have been particularly useful in analyses of both main myzozoan groups, apicomplexans and dinoflagellates: sister groups such as perkinsids, chrompodellids, *Digyalum*, and *Platyproteum*, have all shed light on the evolution of major characters (Goggin and Baker, 1993; Moore et al., 2008; Cavalier-Smith 2014; Janouškovec et al., 2015, 2019; Mathur et al., 2019). But no equivalent sister group to ciliates is known, and most importantly, no group has unambiguously been shown to be sister to the alveolates as a whole (the next neighbour group, the stramenopiles, is distantly related and shares little in common with alveolates).

A handful of predatory protists have been discussed as possible candidates for deep-branching alveolates, based on sequences from one or a small number of genes (Janouškovec et al., 2013; Tikhonenkov et al., 2014; Park and Simpson, 2015). A six gene phylogeny suggested that *Acavomonas* is the sister taxon to myzozoans and *Colponema* is even deeper branching, perhaps as a sister lineage to all other alveolates (Janouškovec et al., 2013). The phylogeny of the small subunit ribosomal RNA gene (SSU rDNA) further showed that *Palustrimonas* and five clades of environmental sequences represent additional deep alveolate lineages (Janouškovec et al., 2013; Park and Simpson, 2015). Their relationships with *Acavomonas, Colponema*, ciliates, myzozoans are unresolved on SSU rDNA trees, but they all appear evolutionarily distinct (Tikhonenkov et al., 2014; Park and Simpson, 2015). Such diversity of deep-branching taxa could be instrumental in interpreting the alveolate origin and early evolution. In particular, an early-branching position for *Colponema* could be consistent with it possessing structures and behaviors thought to be plesiomorphic in the group (Mignot and Brugerolle 1975; O’Kelly, 1993; Tikhonenkov et al., 2014).

Here we report the phylogenomic analyses of transcriptomes of three colponemid lineages, which unambiguously support their monophyly in the phylum Colponemidia, and support their position as the sister lineage to all other alveolates. We further analyse the environmental diversity of colponemids and discuss their role in the ecosystems and importance in understanding the evolution of alveolates.

## 2. Material and methods

### 2.1. Collection, culture establishment and microscopy

Clones Colp-22, Colp-26, and Colp-tractor were obtained from samples collected from freshwater bodies in Ukraine (Seret river plankton; Dniester Canyon; 48°40’01.2”N, 25°51’04.2”E), Con Dao Island (Quang Trung lake; bottom detritus within lotuses; 8°41’31.2”N, 106°36’24.1”E), and British Columbia (City of Delta, Burns Bog; water with plant debris and detritus near a sunken tractor; 49°08’41.8”N 122°55’51.3”W), respectively. Colponemid clone Colp-10 was isolated from desert soil in Morocco (same location and sample as for *Moramonas marocensis* in Strassert et al., 2016); Colponemid clone Colp-15 was isolated from Suoi Da Reservoir in South Vietnam (same location and sample as for *Aquavolon hoantrani* in Bass et al., 2018). Other colponemid strains were obtained as described previously (Janouškovec et al., 2013, Tikhonenkov et al., 2014). Following isolation by a glass micropipette, all novel strains were propagated on the bodonid *Parabodo caudatus* strain BAS-1, which was grown in spring water (Aqua Minerale, PepsiCo or PC Natural Spring Water, President’s Choice) using the bacterium *Pseudomonas fluorescens* as food. Microscopy observations were made using a Zeiss AxioScope A.1 light microscope and JEM-1011 (JEOL) transmission electron microscope as described previously (Tikhonenkov et al., 2014).

### 2.2. DNA and RNA extraction, sequencing, and assembling

Cells of strains Colp-7a, Colp-10, Colp-15, Colp-22, Colp-26, and Colp-tractor were grown in clonal laboratory cultures and were harvested following peak abundance after eating most of the prey. Cells were collected by centrifugation (2000 × *g*, room temperature) on the 0.8 μm membrane of Vivaclear Mini columns (Sartorium Stedim Biotech Gmng, Germany, Cat. No. VK01P042). Total RNA of strains Colp-7a, Colp-10, Colp-15 was extracted using the RNAqueous®-Micro Kit (ThermoFisher Scientific, cat. # AM1931) and was converted into cDNA prior to sequencing using the SMARTer technology (SMARTer Pico PCR cDNA Synthesis Kit, Takara, cat. # 634928). Sequencing libraries were prepared using the Illumina TruSeq protocol and sequenced on an Illumina MiSeq using 250 bp paired end reads. Transcriptome assembly and decontamination were carried out according the pipeline described in Burki et al. (2016). Transcriptomic assemblies are available at NCBI under accession numbers: XXXX–XXXX.

Genomic DNA was extracted from freshly harvested cells using the Epicentre DNA extraction kit (Cat. No. MC85200). SSU rDNA genes of clones Colp-10, Colp-15, Colp-22, Colp-26, and Colp-tractor (GenBank accession numbers: xxxxx) were amplified by PCR using general eukaryotes primers: 18SFU-FAD4 (Colp-10), PF1-FAD4 (Colp-15), EukA-EukB (Colp-22, Colp-26, Colp-tractor) (Medlin et al., 1988; Keeling, 2002; Tikhonenkov et al., 2016). PCR products were subsequently cloned and sequenced (Colp-15) or sequenced directly (Colp-10, Colp-22, Colp-26, Colp-tractor) by Sanger dideoxy sequencing. SSU rDNA genes of other colponemid strains were obtained as described previously (Janouškovec et al., 2013; Tikhonenkov et al., 2014).

### 2.3. SSU rDNA gene phylogeny

The SSU RNA gene phylogeny was based on an earlier dataset (Janouškovec et al., 2013). Sampling was modified to include the new colponemid sequences and a more representative selection of other eukaryotes and to exclude several longer-branching sequences. Sequences were aligned by the localpair algorithm in MAFFT v7.402 (Katoh and Standley, 2013) and trimmed by using the -h 0.4 and -g 0.35 settings in BMGE 1.12 (Criscuolo and Gribaldo, 2010). Maximum likelihood phylogeny was computed in IQ-TREE v1.6.9 (Nguyen et al., 2015) by using the best fit TN+F+R4 model (as determined by the in-built ModelFinder) and 300 non-parametric bootstraps. MrBayes v3.2.6 (Ronquist and Huelsenbeck, 2003) was run with default sampling scheme around the GTR+I+GAMMA4 model space until reaching chain convergence diagnostic (stopval) of 0.01, after which a majority consensus of 2576 trees in the posterior probability distribution was calculated (17,175,000 total generations; burnin 25%).

### 2.4. Phylogenomic dataset construction

The assembled sequences of *Colponema vietnamica* Colp-7a, and colponemid strains Colp-10 and Colp-15 were added to 320 publicly available protein-coding genes from a broad range of taxa (Brown et al., 2018; Strassert et al., 2019) as follows: 1) Protein sequences of the new taxa were clustered with CD-HIT (Fu et al., 2012) using an identity threshold of 85%; 2) Candidates for homologous copies were retrieved by BLASTP searches (Altschul et al., 1990) using the 320 genes as queries (e-value: 1e-20; coverage cutoff: 0.5); 3) In three rounds, phylogenetic trees were constructed and carefully inspected in order to detect and remove paralogs and contaminants and select orthologous copies. For that, sequences were aligned using MAFFT v. 7.310 (Katoh and Standley, 2013) with the -auto flag employed (first round) and MAFFT L-INS-i with default settings (second and third round). Alignments were filtered with trimAL v. 1.4 (Capella-Gutiérrez et al., 2009) using a gap threshold of 0.8 (all three rounds) and single-gene Maximum Likelihood (ML) trees were inferred using FastTree v. 2.1.10 (Price et al., 2010) with the -lg -gamma setting (first round) and RAxML v. 8.2.10 (Stamatakis, 2014) with the PROTGAMMALGF model and 100 rapid bootstrap searches (second and third round).

Seven genes were removed from further analyses because no representative colponemid sequences could be retrieved. Moreover, to allow computationally demanding analyses in later steps, the number of taxa was reduced by excluding several fast-evolving taxa, taxa with more missing data, and taxa outside the ‘TSAR’ clade (Strassert et al., 2019; with exception of a few haptists that were retained as the outgroup). The resulting dataset comprised 313 genes and 71 taxa. Non-homologous characters were removed from individual unaligned sequences with PREQUAL v. 1.01 (Whelan et al., 2018) using a filter threshold of 0.95. Sequences were then aligned with MAFFT G-INS-i using the VSM option (--unalignlevel 0.6) and trimmed with BMGE v. 1.12 (Criscuolo and Gribaldo, 2010) employing -g 0.2, -b 5, -m, and BLOSUM75 parameters. Partial sequences belonging to the same taxon that did not show evidence for paralogy or contamination on the gene trees were merged. The genes (Supplemental Material) were concatenated to a supermatrix with SCaFos v. 1.25 (Roure et al., 2007) and an initial ML tree was calculated using IQ-TREE v. 1.6.9 (Nguyen et al., 2015) with the site-homogeneous model LG+G+F and ultrafast bootstrap approximation (UFBoot, Hoang et al., 2018; 1,000 replicates) employing the -bb and -bnni options. To reduce missing data, based on this tree, some monophyletic strains or species/genera complexes were combined to chimera that were used as operational taxonomic unit (OTU), and a new supermatrix (51 OTUs; 84,451 amino acid positions) was built as described above (see Supplemental Material for the corresponding files).

### 2.5. Phylogenomic analyses

The final supermatrix was used to infer a ML tree using IQ-TREE with the best-fitting site-heterogeneous model LG+C60+G+F with the relatively fast PMSF approach (Wang et al., 2018) to obtain non-parametric bootstrap support (100 replicates). Copies of the obtained tree were manually edited to test alternative topologies using the approximately unbiased test (AU test; Shimodaira, 2002) in IQ-TREE: 1) colponemids sister to all other alveolates (original tree) *versus* colponemids sister to ciliates, 2) colponemids sister to all other alveolates *versus* colponemids sister to myzozoa. Also, the gene alignments that have been used for generating the final supermatrix were subjected to the Multi Species Coalescent (MSC) model in ASTRAL-III (Zhang et al., 2018) to incorporate gene tree uncertainties. ML single-gene trees (Supplemental Material) were inferred using IQ-TREE under the best-fitting substitution model, as determined by the BIC criterion. Branches with bootstrap support <10 were collapsed in the input trees, as recommended by the developers. Quadripartition supports were calculated from quartet frequencies among the set of ML input trees (Sayyari and Mirarab, 2016). Bayesian analyses were conducted using PHYLOBAYES-MPI v. 1.8 (Lartillot et al., 2013) with the CAT+GTR+Γ4 model employing the -dc flag to remove constant sites. Two independent Markov Chain Monte Carlo (MCMC) chains were run for 4,330generations (all sampled). For each chain, the burnin period was estimated by monitoring evolution of the log-likelihood (Lnl) at each sampled point, and 1,000 generations were removed from both chains. A consensus tree of the two chains was built with the bpcomp command and global convergence between chains was assessed by the maxdiff statistics measuring the discrepancy in posterior probabilities (PP). As frequently recognized in PHYLOBAYES, global convergence was not received (for details, see Supplemental Material).

## 3. Results

### 3.1. Phylogenomic analyses

To determine the evolutionary position of colponemids, we generated transcriptomes of *Colponema vietnamica* strain Colp-7a and two newly-discovered relatives, and incorporated their sequences into a phylogenomic dataset (Materials and Methods). All genes were individually inspected for orthology by computing single-gene phylogenies and removing paralogs and contaminant sequences. The final phylogenomic matrix contained 51 OTUs and 313 protein-coding genes (84,451 amino acid positions). Colponemids have high data presence: 99% for the colponemid strain Colp-10, 97% for the colponemid strain Colp-15, and 80% for *C. vietnamica* (see Supplemental Material).

Maximum Likelihood analysis of this dataset recovered a monophyly of Colponemidia, which was placed as a sister lineage to all other alveolates with maximal bootstrap support (Fig. 1), in agreement with Bayesian analysis (Figs 1 and S1). The AU test supports this topology (p-AU = >0.95) by rejecting the alternative hypothesis of a colponemids/ciliates sister relationship at the p-AU = <0.05 significance level. The AU test also rejects the position of colponemids sister to Myzozoa (p-AU = 0; with p-AU = 1 for colponemids sister to all other alveolates). In agreement with previous studies and current systematics, the Maximum Likelihood analysis also recovered monophyletic alveolates, ciliates, myzozoans, core dinoflagellates, and apicomplexans, as well as the sister relationship between alveolates and stramenopiles, each fully supported (Strassert et al., 2019). Bayesian analysis showed the same results (Figs 1 and S1): two Markov chains recovered monophyletic colponemids as a sister group to other alveolates with maximal posterior probability support, as well as monophyletic ciliates, myzozoans, alveolates, stramenopiles, rhizarians, and ‘SAR’.

**Figure 1.**
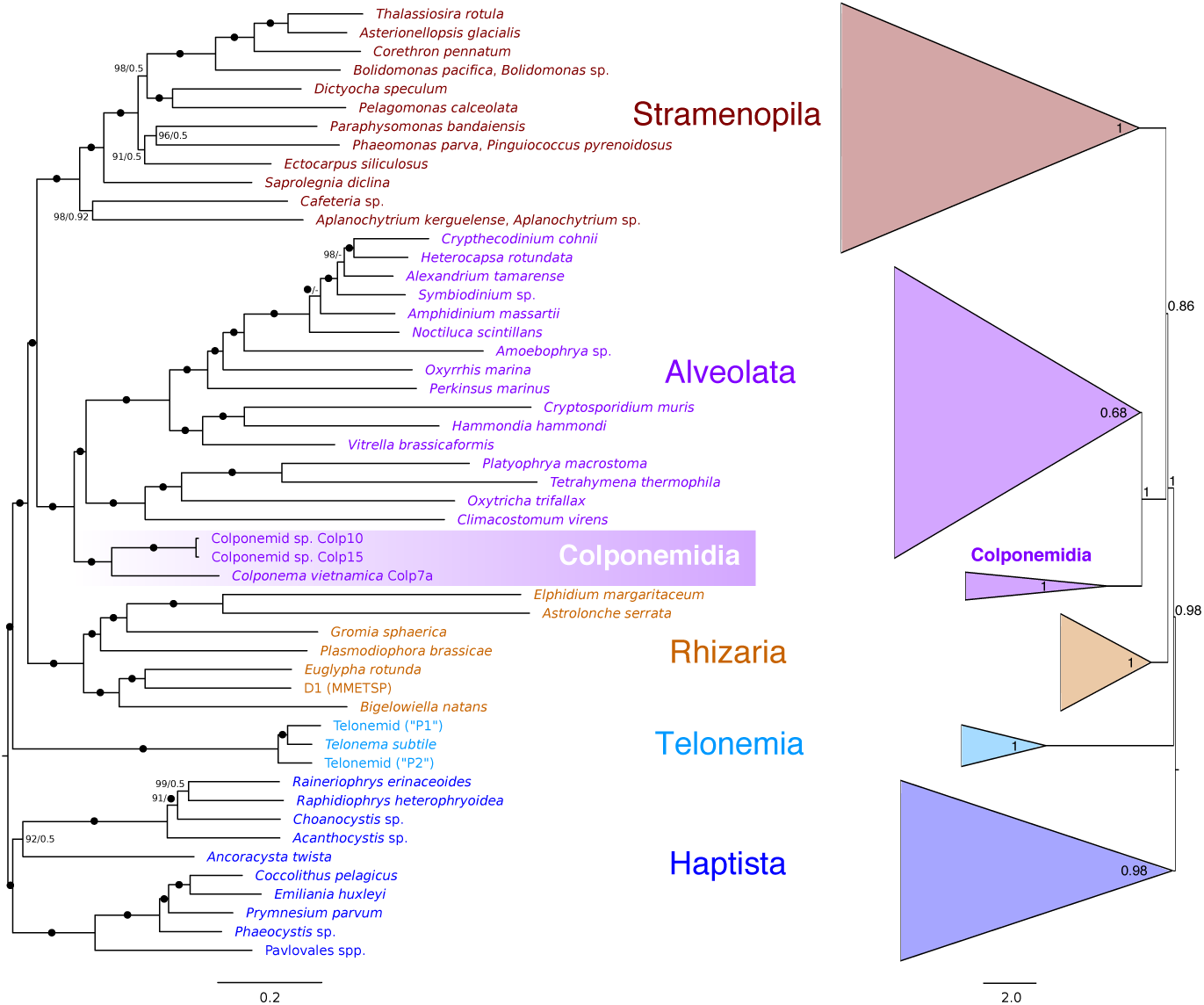
Phylogenomic trees showing the sister group relationship between colponemids and all other alveolates. The tree on the left side represents the best Maximum Likelihood (ML) tree inferred from a concatenated alignment of 313 protein-coding genes using the LG+C60+G+F-PMSF model. Numbers at nodes indicate bootstrap support (left) and Bayesian posterior probability (right). Full support in both analyses is shown by black circles. Maximum support in only one of the two analyses is indicated by a black circle appended to a number. Dashes at nodes indicate topological incongruence with the Phylobayes tree (see Figure S1). The tree on the right is based on summary-coalescent analyses (Astral-III) of 313 single-gene ML trees (Supplemental Material) using the same taxa as shown on the left (collapsed to polygons). The numbers at nodes are local posterior probabilities.

The result of the multi species coalescent analysis is in an overall agreement with the Maximum Likelihood topology (Fig. 1). It also indicates a sister relationship of colponemids to all other alveolates although only with modest local posterior probability support.

### 3.2. Diversity, distribution, and ecological role of Colponemidia

We previously isolated and described several strains of *C. edaphicum* and *C. vietnamica* from different freshwater and soil environments of Chukotka, Caucasus, and Vietnam (Mylnikov, Tikhonenkov, 2007; Janouškovec et al., 2013; Tikhonenkov et al., 2014). The two new colponemid strains isolated here from Morocco (Colp-10) and Vietnam (Colp-15) are considerably divergent from both species (Figs 1 and 2a). Additionally, we have isolated three new colponemid strains (Colp-22, Colp-26, and Colp-tractor) from a river in Ukraine, lake in South Vietnam, and bog in British Columbia, Canada. All the three strains share the overall *C. vietnamica* morphotype (e.g., they have elongated-oval not flattened cells with comparatively short ventral groove and anterior flagellum as well as long posterior flagellum, which sometimes undulates in the longitudinal ventral groove), and a phylogenetic analysis of their SSU rRNA genes confirms that they belong to the *C. vietnamica*/*edaphicum* clade (Figs 2a and 3).

**Figure 2.**
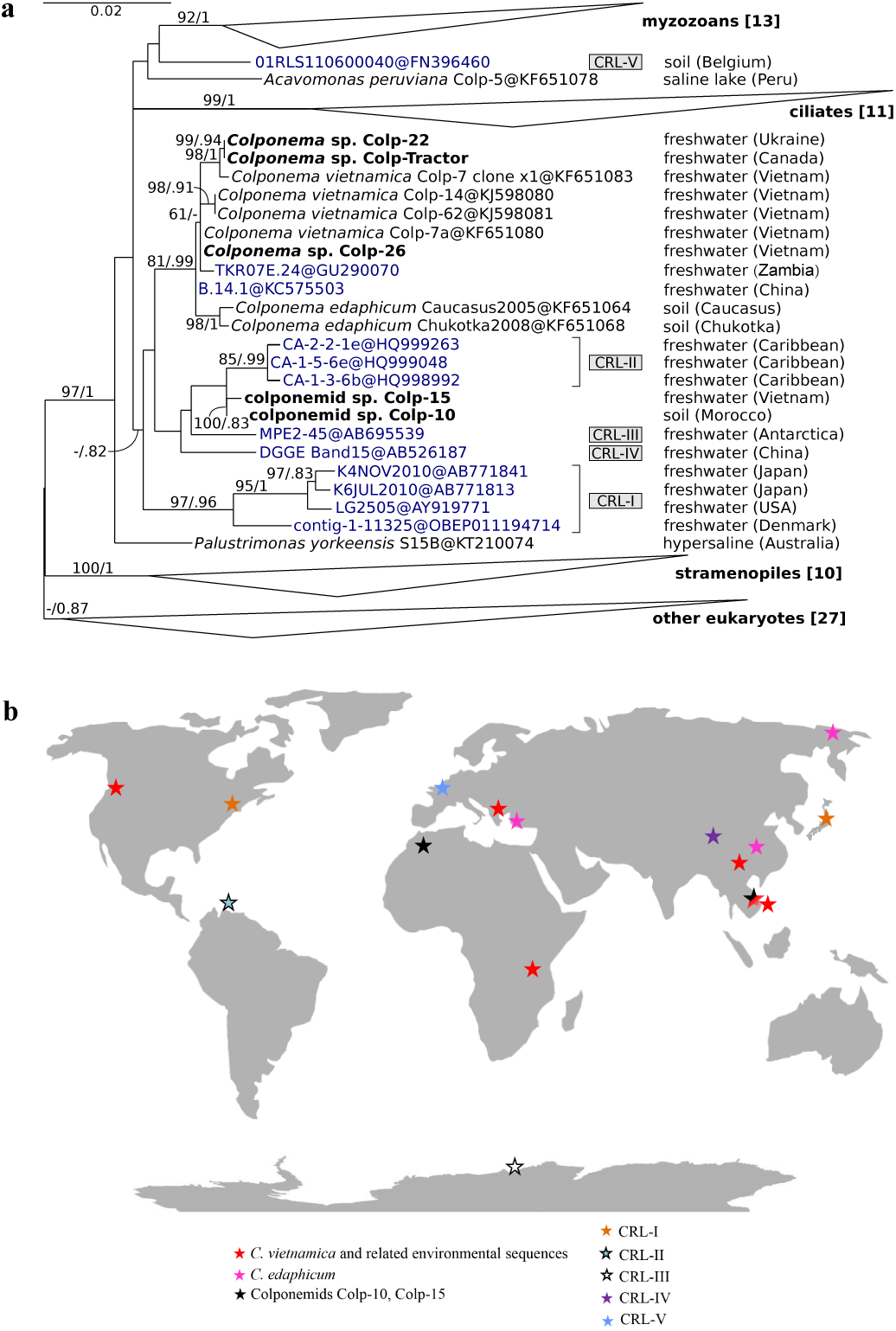
The small subunit ribosomal RNA gene phylogeny (**a**) and geographic distribution (**b**) of colponemids and environmental colponemid-related lineages (CRLs). A maximum likelihood tree of 86 sequences (best fit TN+F+R4 model) is shown with non-parametric bootstrap supports and Bayesian posterior probabilities at branches (values >50/>0.7 are shown). Environmental sequences are highlighted in dark blue and include five previously identified colponemid-related lineages (CRLs; grey boxes). Sequence accession numbers in GenBank are shown after the @ sign. Numbers of species in compacted clades are shown in square brackets.

**Figure 3.**
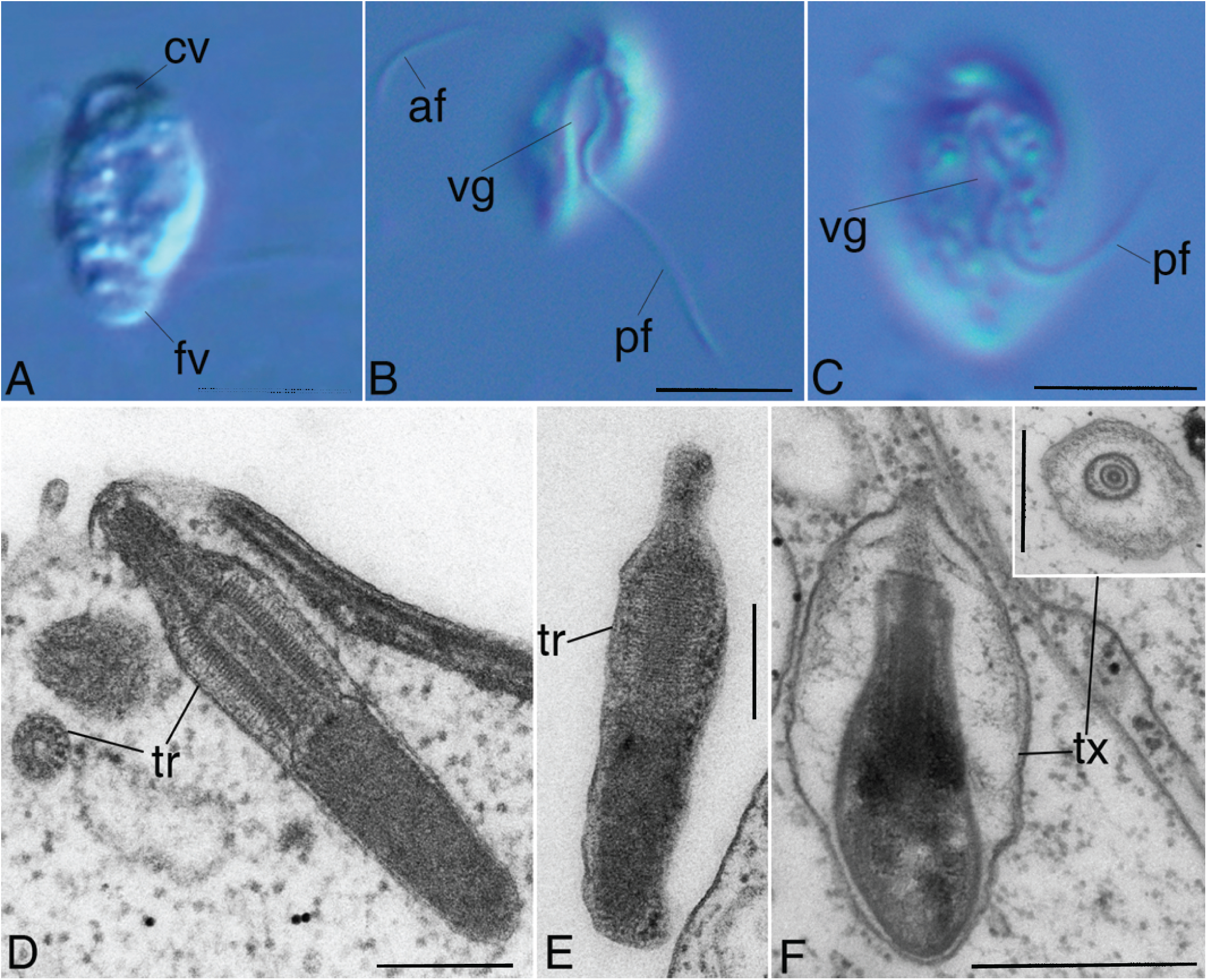
External morphology and extrusive organelles of colponemids. **A–C** – general view of *C. vietnamica* (differential interference contrast) showing heterodynamic anterior (af) and posterior (pf) flagella, pronounced ventral groove (vg) with undulating posterior flagellum, apical contractile vacuole (cv), and posterior food vacuole (fv). **D, E** – Ultrathin sections (TEM) of skittle-shaped trichocysts (tr) with an extended tip, roundish in a cross-section of colponemid clone Colp-10. **F** – Longitudinal and cross (inset) sections (TEM) of bottle-shaped toxicyst (tx) enclosed inside a vesicle of *Colponema vietnamica.* Scale bars: A–C – 5 μm, D, E – 200 nm, F – 500 nm.

We examined SSU rDNA sequences from colponemid-related lineages (CRLs) from environmental surveys retrieved from GenBank (Fig. 2a), which confirmed colponemids are not common in natural surveys, but are diverse and widespread, and also potentially form multiple subgroups basal to alveolates. Within the main colponemid clade, two environmental sequences are closely related to *C. edaphicum, C. vietnamica*, and three new strains (Colp-22, Colp-26, and Colp-Tractor). Five other environmental sequences previously referred to as CRL-II, III, IV (Janouškovec et al., 2013) form a weakly-supported subgroup with Colp-10 and Colp-15. Maximum Likelihood and Bayesian trees also show two groups of environmental sequences, CRL-V and CRL-I, to be independent of other colponemids, but again without statistical support (Fig. 2a).

Geographical distribution analysis has shown that colponemids are cosmopolitan at the species level (*C. vietnamica*) (Fig. 2b). They have been observed in all hemispheres, and in different geographic zones including the tropical, subtropical, temperate, and both Arctic and Antarctic polar zones. Interestingly, all molecularly identified colponemids and CRLs sequences have never been found in marine samples, and are only known to inhabit freshwater ecosystems and soil (i.e., thin freshwater films surrounding soil particles). They are present in different types of biotopes including plankton, bottom detritus, plant debris, peatlands, aquatic mosses, and wastewater from treatment facilities, but they are not known as an abundant component of microbial communities.

All newly studied colponemids are obligate eukaryovores and feed on other small protists in all observations from natural samples and in laboratory cultures (e.g., on bodonids and chrysophytes), which is consistent with the presence of prey-immobilizing toxicysts in microscopically investigated species *Colponema loxodes, C.* aff. *loxodes, C. edaphicum*, and *C. vietnamica* (Mignot, Brugerolle, 1975; Myl’nikova and Myl’nikov, 2010; Tikhonenkov et al., 2012, 2014). However, many dinoflagellates, colpodellids, and *Psammosa* possess trichocysts (Rosati, Modeo, 2003; Okamoto et al., 2012) but not toxicysts, while ciliates have both. Interestingly, the cells of colponemid strain Colp-10 have elongated skittle-shaped trichocysts (∼0.9 × 0.15 μm) with an extended tip, roundish in a cross-section (Fig. 3). The presence of both types of extrusive organelles in a single alveolate phylum, the Colponemidia, is notable and increases our understanding of early alveolate morphology.

## 4. Discussion

### 4.1. Phylogeny and diversity of Colponemidia

Previous phylogenetic analyses of *Colponema* based on rDNA and a set of six genes failed to conclusively resolve its evolutionary position. SSU and SSU/LSU rDNA phylogenies placed it as a sister to myzozoans, or myzozoans+*Acavomonas* (Janouškovec et al., 2013, 2017b; Mikhailov et al., 2014; Tikhonenkov et al., 2014; Cavalier-Smith, 2018), or sister to myzozoans+*Acavomonas*+*Palustrimonas* (Park and Simpson, 2015). Hsp90 and alpha tubulin phylogenies placed *Colponema* as a sister to ciliates, whereas concatenations of SSU and LSU with combinations of HSP90, actin, and alpha- and beta-tubulin placed it basal to other alveolates without statistical support (Janouškovec et al., 2013). Our phylogenomic analyses of 313 eukaryote protein-coding genes from several colponemids now resolves this position, unambiguously placing them as a sister to all other alveolates with strong support in Maximum Likelihood site-heterogeneous mixture model and Phylobayes estimations.

Colponemids are widespread in nature and found in a broad range of habitats, including plankton, bottom sediments, boggy sites, and soil, and in extreme environments such as Arctic permafrost (*C. edaphicum*) and desert soil (colponemid Colp-10). Their widespread, or perhaps cosmopolitan nature is common in small heterotrophic flagellates in general (Lee, Patterson, 1998; Azovsky et al., 2016), which are also the primary food resources for colponemids. Exactly how they survive hostile conditions remains to be determined; only Colp-10 possesses cysts in its life cycle, and the survivability of *C. edaphicum* in 28–32 millennial permafrost (Shatilovich et al., 2009, 2010) remains a mystery. Despite this, however, all molecularly-identified colponemids to date originate from freshwaters and freshwater soil microhabitats — never from marine habitats.

### 4.2. Colponemids and the evolution of alveolates

Taking the structural features of colponemids into consideration together with their basal position in the tree has a major impact on how we reconstruct the common ancestor of alveolates and the major sub-groups. In particular, the overall picture of the ancestral alveolate that emerges is somewhat different from the characteristics that typify the three major alveolate groups individually: it was a free-living predatory protist with submembrane alveolar vesicles, two heterodynamic flagella, tubular mitochondrial cristae, a posterior digestive vacuole, and extrusive organelles (trichocysts or toxicysts) for active hunting by phagocytosis. This description is a close match with modern colponemids, and indeed they have been described as “living fossils” (Cavalier-Smith, 2018). Interestingly, however, the morphostasis of colponemids may go even deeper, because they also resemble excavate protists (O’Kelly, 1993), with ventral groove, tubular mitochondrial cristae, and vane on the posterior flagellum (although the vane is ventral in *Colponema* and dorsal in excavates).

Fine structure of the colponemid cytoskeleton needs further investigation, as even here colponemids appear to retain ancient characteristics (Tikhonenkov et al., 2014). In the flagellar root system, the so-called R2 fibre splits into oR2 and iR2 that support the feeding apparatus, and this split likely represents an ancient characteristic of the last common ancestor of excavates, stramenopiles, apusozoans, amoebozoans, collodictyonids, haptophytes, cryptophytes, and probably alveolates (Yubuki, Leander, 2013). The cytoskeleton configuration in *Colponema* would therefore be ancestral not just to alveolates but to all eukaryotes (Tikhonenkov et al., 2014).

Cytoskeletal elements for feeding and locomotion are connected functionally and historically. In bacteriotrophs (most heterotrophic protists), flagellar beating creates a current directed to the cytostome by the feeding groove, moving suspended bacteria to the ‘mouth’. Microtubular bands originating at the basal body provide mechanical rigidity to the groove margins. Colponemids retain this ancestral organization, although they phagocytize eukaryotes and not bacteria, and use active hunting of mobile prey rather than passive suspension feeding. Subsequently, the ancestor or myzozoans and *Acavomonas* probably lost the link between flagellar apparatus and feeding groove as it transitioned from bacteriotrophy and phagotrophy to suction feeding on larger cells. For example, *Acavomonas* has no pronounced ventral groove (Tikhonenkov et al., 2014). But the newly-developed feeding system based on the apical complex retained its link to the basal bodies because the microtubular band formerly supporting the feeding grove was most likely converted into the conoid that strengthens the rostrum and the edge of the cytostome in extant myzozoans. This link is obscured by the complex life cycles of most myzozoans, but can be clearly observed in the direct link between basal bodies and conoid in *Psammosa* (Okamoto, Keeling, 2014).

The origin of myzocytosis and radiation of myzocytosis-feeding taxa may have been response to the increase of eukaryote (prey) cells sizes. Macrophagous predatory protists select prey by size, at that larger potential prey is more difficult to swallow (Bengtson, 2002). Unicellular eukaryotes reached their maximum size before the Cambrian explosion, although an increase in the average cell size through the Proterozoic was not a continuous trend (Huntley et al., 2006). The observed size ratios between modern predatory protists (dinoflagellates, chrysophytes, bicosoecids, pedinellids, ciliates) and their prey range from 0.4:1 to 30:1 because of different mechanisms of prey capture (Hansen et al., 1994). Myzocytosis allows a predator to consume biomass accumulated in prey of any size, for example, *Colpodella gonderi* attacks large ciliates (Olmo at al., 2011). The ancestors of dinoflagellates and apicomplexans were likely capable of myzocytosis and possessed an apical complex-like structure including an open conoid, rhoptries, and micronemes. These characteristics are mostly lost in autotrophic dinoflagellates and their relatives, but parasitic apicomplexans and dinoflagellates have adapted their myzocytotic feeding to multicellular organisms. Although parasitic dinoflagellates and perkinseids possess flagellar apparatus to search their multicellular or unicellular prey, apicomplexan parasites have lost flagellated zoospores. This loss could be associated with the events in the Middle Cambrian, known as the “Agronomical revolution”, or the “Cambrian substrate revolution” (Seilacher and Pflüger, 1994). At this time, organic material began to accumulate on the seabed and burrowing organisms increased in number and diversity. The immobile resting oocysts of apicomplexans have got high chances of being acquired by the burrowing host. Further evolution without mobile infectious stages has led to complex apicomplexan life cycles exploiting the behavior of animal vectors.

Another well-studied character in alveolate evolution is the plastid organelle. Photosynthetic and non-photosynthetic plastids derived from a secondary enslavement of a red alga are found throughout the myzozoans (Janouškovec et al., 2015), but no evidence for plastids or plastid genes has been found in colponemids to date (Janouškovec et al., 2013). The origin of this plastid, has been widely debated: on one side it is argued to be the result of a single ancient plastid origin predating alveolates, stramenopiles and perhaps several other lineages (Cavalier-Smith, 1999, 2013), while on the other side it has been argued to have originated more recently within alveolates by more hierarchical endosymbioses (Petersen et al., 2014; Stiller et al., 2014). If the former is true, then the ancestor of alveolates would have been mixotrophic and colponemids (and ciliates) lost plastids and plastid genes secondarily (as have parasitic myzozoans *Cryptosporidium* and *Hematodinium*: Gornik et al., 2015; Xu et al., 2004; Zhu et al., 2000). In contrast, the latter possibility suggests colponemids never had plastids and are more similar to the ancestral host of myzozoan plastids. Distinguishing between the heterotrophic *versus* mixotrophic nature of the ancestral alveolate has direct ramifications for the origin of plastids in the photosynthetic stramenopiles, haptophytes and cryptophytes, and plastid presence or loss in the related heterotrophs. As such, the origin of alveolate plastids could resolve a major gap in our understanding of the evolution of photosynthesis in a major part of eukaryotic diversity and uncover the general principles governing plastid acquisition and loss.

## Acknowledgements

We thank A.A. Abramov, A.V. Shatilovich, Y.V. Dubrovsky, L. Nguyen-Ngoc, H. Doan-Nhu, E. S. Gusev, A.N. Tsyganov, and the staff of the Russian-Vietnam Tropical Centre, Coastal Branch, Nha Trang, especially Hoan Q. Tran, Tran Duc Dien, and Nguyen Thị Hai Thanh for assistance with sample collection and trip management. Field work in Vietnam is part of the project ‘Ecolan 3.2’ of the Russian-Vietnam Tropical Centre. This work was supported by grants from the Russian Foundation for Basic Research (grant nos. 17-04-00899 and 18-504-51028), the Ministry of Education and Science of the Russian Federation (project No. AAAA-A18-118012690098-5), and from the Natural Sciences and Engineering Research Council of Canada (NSERC) (grant no. 227301).

## Supplementary material

**Figure S1.** Phylobayes trees inferred from a concatenated alignment of 313 protein-coding genes under the CAT+GTR+Γ4 model. For more information, please see text next to the trees.

**OTU_list.txt.** Text file listing the taxa used in this study and their completeness in the supermatrix. Note, several strains or species/genera complexes were combined to chimera that were used as operational taxonomic unit (OTU).

**Supermatrix.fasta.** Concatenated trimmed alignment of 313 protein-coding genes.

**Single_gene_alignments.** Directory containing all single-genes alignments.

**Single_gene_ML_trees.tre.** Tree file with 313 ML single-gene trees.

---

**Figure S1a.**
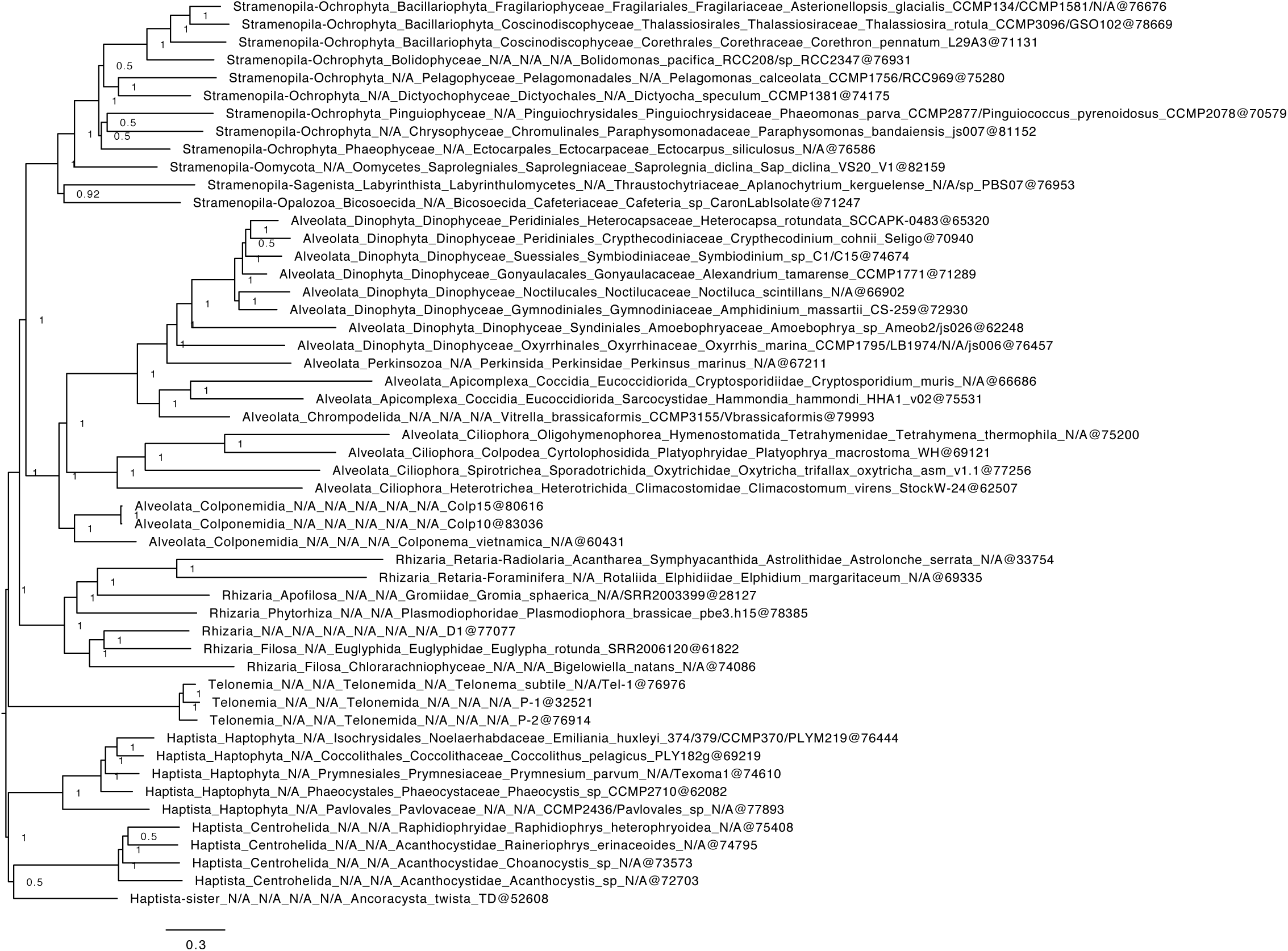
Phylobayes tree inferred from a concatenated alignment of 313 protein-coding genes under the CAT+GTR+Γ4 model. The tree is a consensus from two MCMC chains (Figure S1b and S1c) that were run for 4,330 generations with a burnin of 1,000 generations. Node support is given by Bayesian posterior probabilities.

**Figure S1b.**
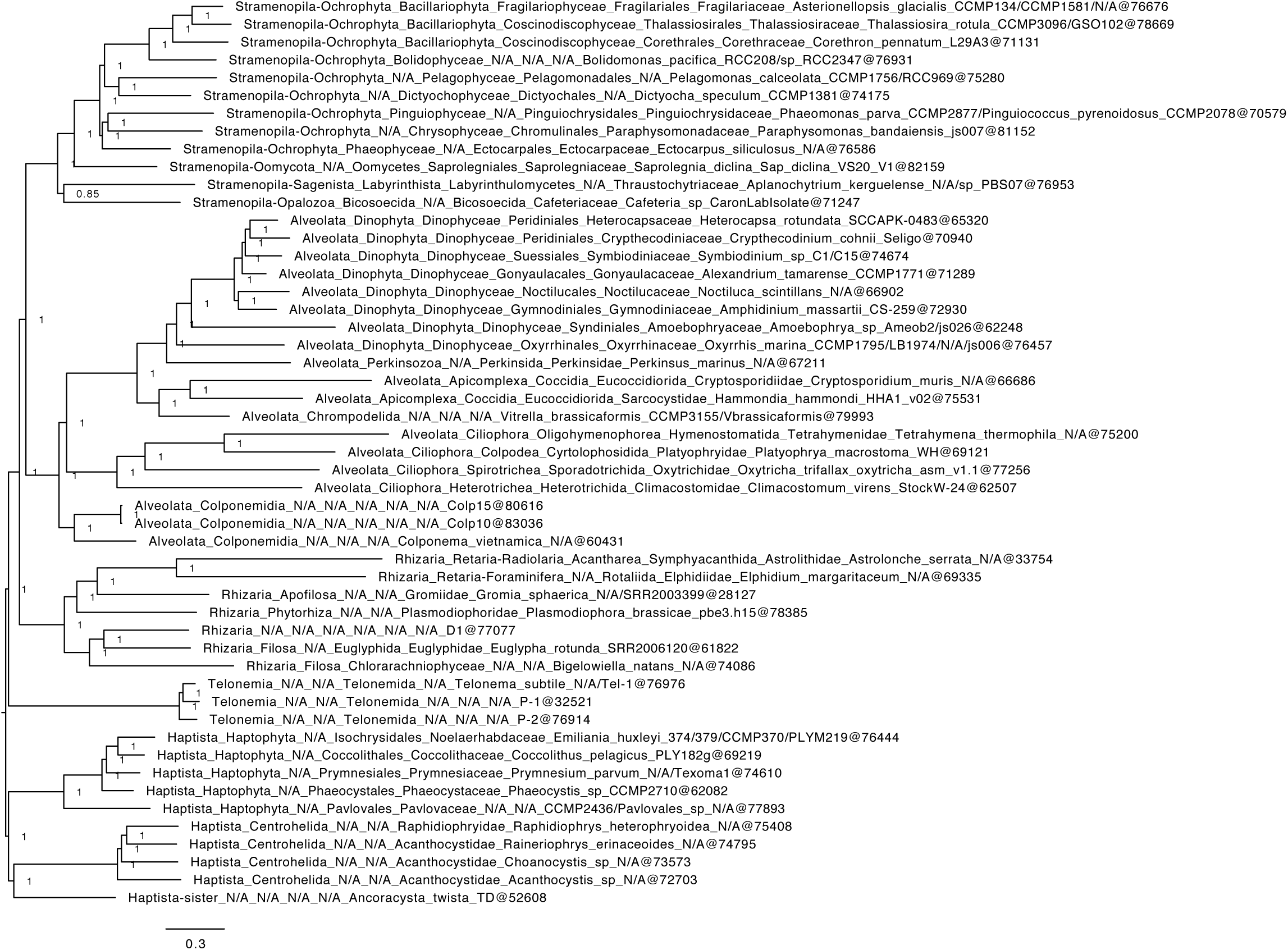
Phylobayes tree inferred from a concatenated alignment of 313 protein-coding genes under the CAT+GTR+Γ4 model. The tree is a consensus from the first MCMC chain used in Figure S1a. Node support is given by Bayesian posterior probabilities.

**Figure S1c.**
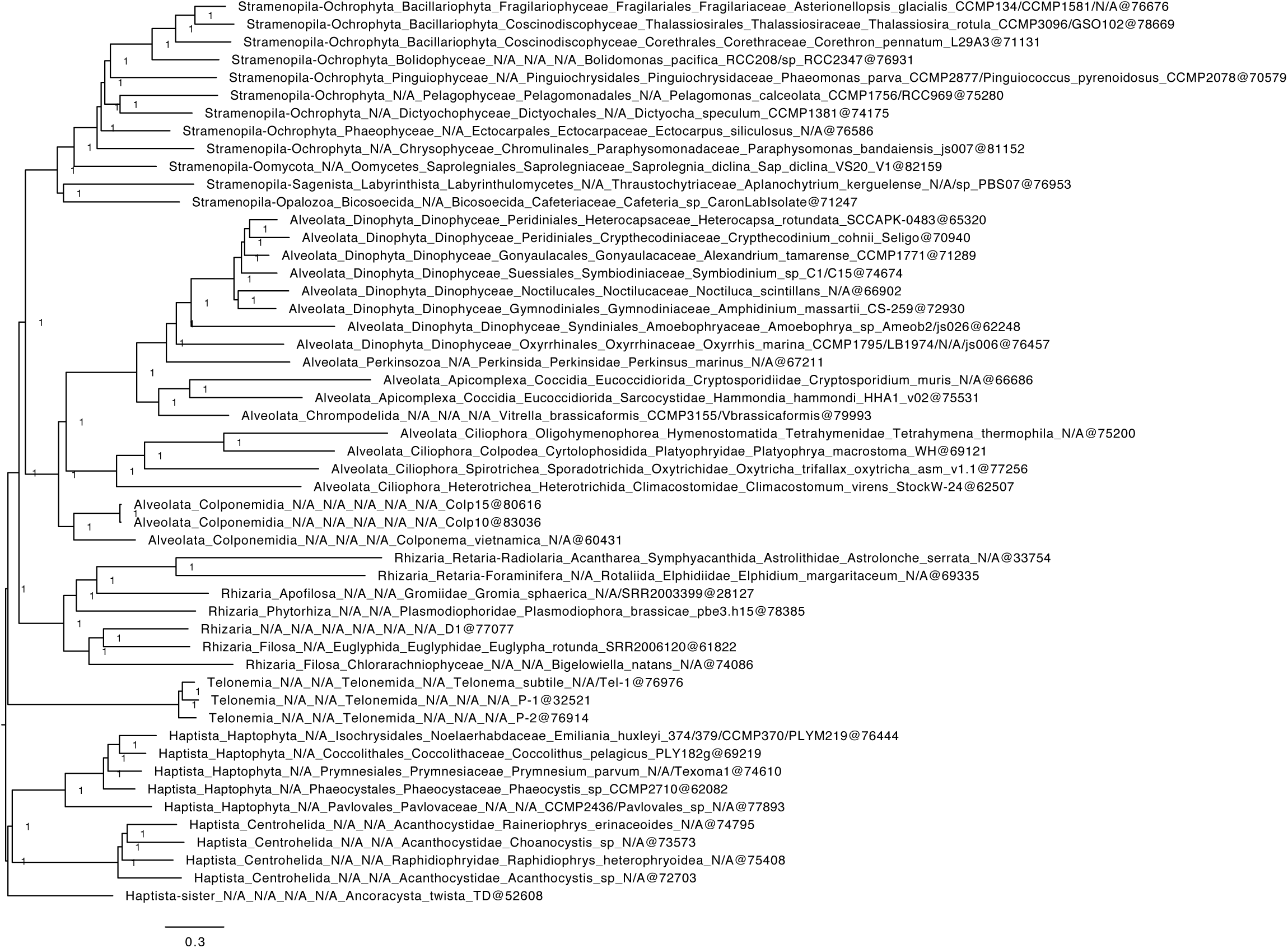
Phylobayes tree inferred from a concatenated alignment of 313 protein-coding genes under the CAT+GTR+Γ4 model. The tree is a consensus from the second MCMC chain used in Figure S1a. Node support is given by Bayesian posterior probabilities.

